# Gating mechanism of the human α1β GlyR by glycine

**DOI:** 10.1101/2023.08.08.552474

**Authors:** Xiaofen Liu, Weiwei Wang

## Abstract

Glycine receptors (GlyRs) are members of the Cys-loop receptors that constitute a major portion of neurotransmitter receptors in the human nervous system. GlyRs are found in the spinal cord and brain mediating locomotive, sensory and cognitive functions, and are targets for pharmaceutical development. GlyRs share a general gating scheme with Cys-loop receptor family members, but the underlying mechanism is unclear. Recent resolution of heteromeric GlyRs structures in multiple functional states identified an invariable 4:1 α:β subunit stoichiometry and provided snapshots in the gating cycle, challenging previous beliefs and raising the fundamental questions of how α and β subunit functions in glycine binding and channel activation. In addition, how a single glycine-bound extracellular domain conformation leads to structurally and functionally different open and desensitized states remained enigmatic. In this study, we characterized in detail equilibrium properties as well as the transition kinetics between functional states. We show that while all allosteric sites bind cooperatively to glycine, occupation of 2 sites at the α-α interfaces is necessary and sufficient for GlyR activation. We also demonstrate differential glycine concentration dependence of desensitization rate, extent, and its recovery, which suggests separate but concerted roles of ligand-binding and ionophore reorganization. Based on these observations and available structural information, we developed a comprehensive quantitative gating model that accurately predicts both equilibrium and kinetical properties throughout glycine gating cycle. This model likely applies generally to the Cys-loop receptor family and informs on pharmaceutical endeavors in function modulation of this receptor family.

## Introduction

Cys-loop receptors are pentameric ligand-gated ion channels that constitute a major portion of ionotropic neurotransmitter receptors in the mammalian nervous systems, including the cation-selective nicotinic acetylcholine receptors (nAChR) and type 3 serotonin receptors (5-HT3), as well as the anion-selective glycine receptors (GlyR) and GABA(A) receptors(1, 2). GlyRs mediate both locomotive and sensory signals in the spinal cord, whose disfunction causes hyperekplexia (startle syndrome)(3, 4) and relates to chronic inflammatory pain(5, 6), making them potential drug targets(7-10). Autism spectrum disorders are also genetically associated with defects of GlyRs, implicating their indispensable function in the brain(11, 12).

GlyRs share a general gating scheme with other Cys-loop receptors with unclear underlying mechanism. Upon binding with glycine, GlyRs quickly transition from non-conductive resting state to conductive open state. The open state then turns into non-conductive desensitized state, and only recovers by the removal of glycine. In early developmental stages GlyRs are homomeric containing 5 α subunits, while in adult animals they are heteromeric consisting of 4 α and 1 β subunits (13-16). Recent heteromeric GlyR structures contained both fully and partially occupied glycine binding sites, all of which share an identical pseudo-symmetric extracellular glycine-binding domain (ECD) conformation, and with no clear correlation with the open or desensitized states(14-16). This raises the question of how many glycine bindings are required, and whether different subunit types contribute differentially for GlyR activation, like other Cys-loop receptors(17-22). In addition, how the same glycine bound ECD conformation results in different functional states (open and desensitized) is puzzling.

To address the above questions, we systematically characterized the kinetical and thermodynamical properties of the wild-type human α1β GlyR and its mutants throughout the glycine gating cycle. We show that 2 allosteric sites at the α-α interfaces are necessary and sufficient for GlyR activation, while sites at the β subunits contribute through binding cooperativity. We also show simple mass action-based behavior for open and close, but glycine concentration-independent desensitization rate, suggesting that the open to desensitization transition being purely within the transmembrane pore-forming domain (TM). The more extensive desensitization and slower recovery from higher glycine concentrations can be attributed to slower glycine binding/dissociation kinetics in the activated ECD conformation. These observations were consolidated into a mathematical model that encompasses all GlyR gating parameters and closely predicts channel behavior. Together, this work provides a quantitative description that addresses fundamental question regarding GlyR gating mechanism, and reconciliate the seemingly discrepancies in structural and functional studies.

## Results

### 2 α1-α1 allosteric sites are sufficient and necessary for activation

Recent heteromeric glycine receptor structures revealed that all the 5 ECD allosteric sites in each pentameric channel are capable of glycine binding(14-16) (Fig. 1A, B). Although less than full occupancy seemed sufficient for activation(16), the exact number was unknown. In addition, whether all three types of binding sites (Fig. 1B), α(+)-α(-), β(+)-α(-) and α(+)-β(-), share identical contribution to channel activation is unclear. Since heteromeric GlyRs are found to have an invariable stoichiometry of 4α:1β(14-16), strategic allosteric site mutations in α and/or β subunits, as well as in their concatemers allowed for the evaluation of function of individual sites. For this, we generated multiple α1-β and α1-β-α1 concatemeric subunits and identified the ones that retained wild-type glycine activation profile for further mutagenesis (Supplementary Fig. 1). Combinations of (+) and (-) side mutations at the α1, β subunits, as well as α1-β and α1-β-α1 concatemers enumerated a panel of α1β GlyRs containing 1∼5 intact glycine binding sites (Fig. 1C, Table 1). In one mutant, α1:α1-β-α1 FYF GlyR, 2 intact α1-α1 sites were sufficient for glycine activation, showing similar maximum current as wild-type (4.5 ± 0.7 nA mutant v.s. 7.0 ± 0.9 nA wild-type, Fig. 1C, D and Table 1) and slightly increased EC_50_ from ∼100 μM to ∼240 μM. Removing one more intact site (leaving only 1) as in α1:α1-β-α1 RFYF rendered the channel inactive, while adding more sites (3 and 4 intact sites total) slightly decreased EC_50_. These data demonstrates that a minimal of 2 intact sites are sufficient for heteromeric GlyR activation. We further tested whether allosteric sites at α1-α1, α1-β or β-α1 subunit interfaces have same contribution to activation (Fig. 1E, F and Table 1). As expected, only 1 intact site at the β-α1 interface was insufficient for activation. Moreover, unlike at α1-α1, 2 intact sites at α1-β and β-α1 interfaces did not activate GlyR. Even the addition of one more α1-α1 site (3 in total) did not result in active GlyR. These recordings suggest that 2 α1-α1 allosteric sites are both sufficient and necessary for activation, while α1-β or β-α1 sites are neither sufficient nor necessary, but only decrease EC_50_. These findings are consistent with available structural data where activated states (open/desensitized) are always found with not less than 2 glycine bound in allosteric sites at the α-α interfaces.

**Figure 1.**
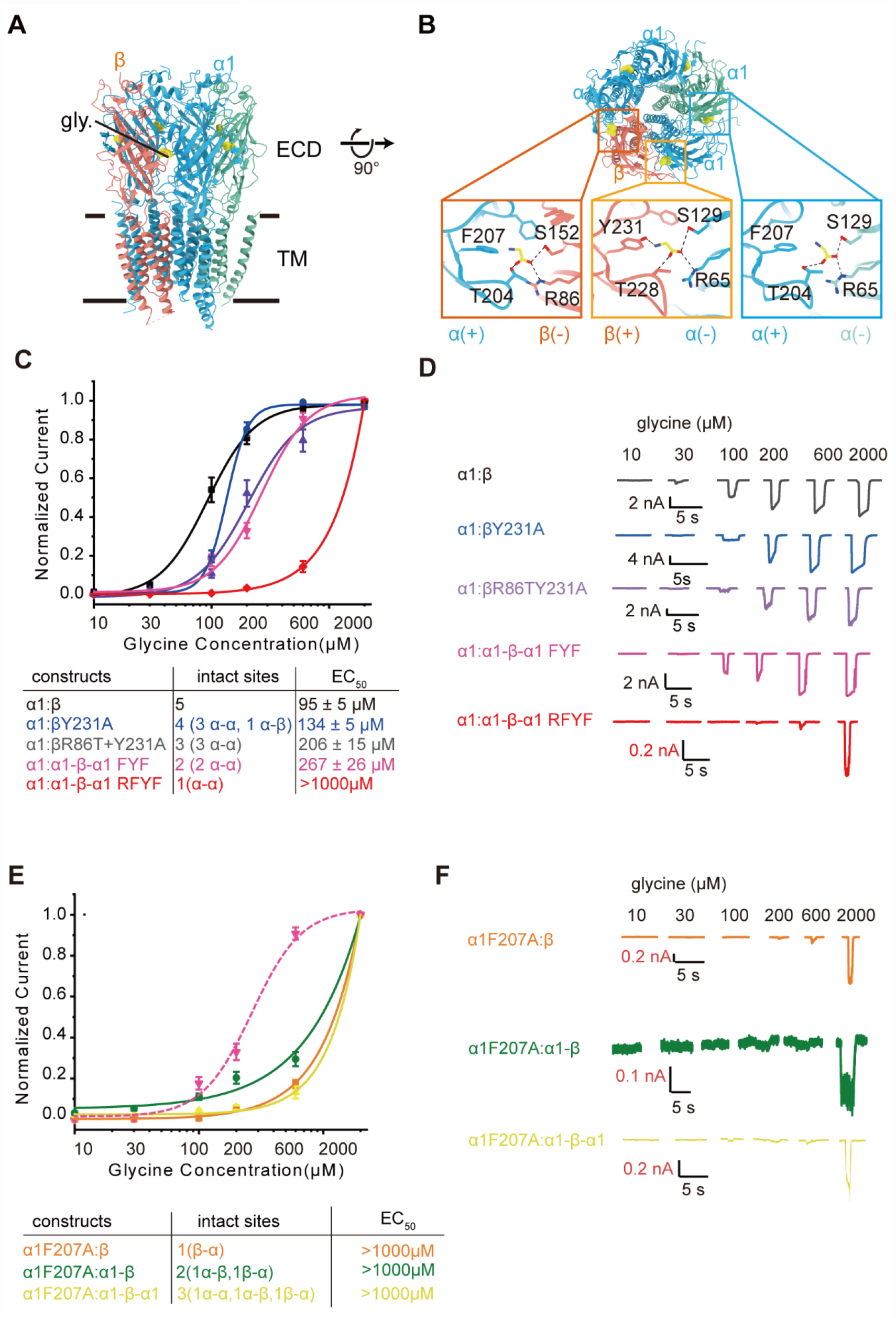
Glycine response of α1β GlyR with mutations in specific allosteric sites. 2 allosteric sites at the α-α subunit interface are sufficient and necessary for activation. (A) Side view and (B) top-down view of the α1β GlyR atomic models in complex with glycine. Glycine (yellow) binding at 3 types of pockets at the (+) and (-) sides of the α (sky blue/green) and β subunits (salmon) are shown with key amino acid residues. (C) Glycine dose-response of wild-type (α1:β) and indicated mutants (α1: α1-β-α1 FYF denotes α1:α1F207A-βY231A-α1F207A and α1: α1-β-α1 RFYF denotes α1: α1R65DF207A-βY231A-α1F207A. n = 8 - 15, mean ± S.E.M.). (D) Representative whole-cell recordings for derivation of dose-response curves shown in C. (E) Glycine dose-response of mutants containing less than 2 intact α1-α1sites (n = 7 - 10, mean ± S.E.M.). α1: α1- β-α1 FYF (hot pink dashed line) is shown for comparison. (F) Representative whole-cell recordings for E. Responses are normalized to max response at 1 mM glycine. The number and types of intact sites, together with EC_50_s are shown in the table below.

**Table 1.** Hill fit parameters of glycine dose-response of wild-type and mutant GlyRs in figure1.

| Constrcuts | EC<sub>50</sub> ( μM) | $n_h$(Hill slope) | $n$ (Cell) | $I_{max}$ (nA) |
| --- | --- | --- | --- | --- |
| $\alpha 1:\beta$ | 95 ± 5 | 2.5 ± 0.4 | 12 | 7.0 ± 0.9 |
| $\alpha 1:\beta Y231A$ | 134 ± 5 | 4.7 ± 0.6 | 15 | 7.5 ± 1.0 |
| $\alpha 1:\beta R86TY231A$ | 206 ± 15 | 2.1 ± 0.3 | 14 | 7.1 ± 0.5 |
| $\alpha 1:\alpha 1-\beta-\alpha 1$ FYF | 267 ± 26 | 2.2 ± 0.4 | 8 | 4.5 ± 0.7 |
| $\alpha 1:\alpha 1-\beta-\alpha 1$ RFYF | > 1000 | 1.6 ± 2.0 | 8 | 0.8 ± 0.3 |
| $\alpha 1F207A:\beta$ | >1000 | 1.4 ± 1.5 | 10 | 0.8 ± 0.2 |
| $\alpha 1F207A:\alpha 1-\beta$ | > 1000 | 1.0 ± 0.6 | 8 | 0.2 ± 0.1 |
| $\alpha 1F207A:\alpha 1-\beta-\alpha 1$ | > 1000 | 1.8 ± 3.3 | 7 | 0.5 ± 0.1 |

EC<sub>50</sub>: medium effective concentration of glycine.
$I_{max}$ : maximum glycine-induced current amplitude.

**Table 2.** Summary of channel kinetical and thermodynamical properties during glycine activation.

| | Glycine(μM) | $T_{Open}$(s) | $T_{Close}$(s) | $T_{des}$(s) | $T_{des}$(s) | $I_{des} / I_{max}$ | $n$(cells) |
| --- | --- | --- | --- | --- | --- | --- | --- |
| $\alpha 1wt\beta wt$ | 40 | $0.6 \pm 0.1$ | $0.3 \pm 0.1$ | $4.8 \pm 0.9$ | $7.7 \pm 0.6$ | $0.66 \pm 0.04$ | 5 |
| | 100 | $0.18 \pm 0.03$ | $0.3 \pm 0.1$ | $7.5 \pm 1.2$ | $15.7 \pm 0.6$ | $0.39 \pm 0.04$ | 5 |
| | 200 | $0.07 \pm 0.01$ | $1.1 \pm 0.1$ | $7.8 \pm 0.6$ | | $0.23 \pm 0.03$ | 8 |
| | 1000 | $0.06 \pm 0.01$ | $1.0 \pm 0.1$ | $6.5 \pm 0.4$ | $57 \pm 8$ | $0.18 \pm 0.02$ | 13 |
| $\alpha 1s\beta sg$ | 40 | $0.9 \pm 0.1$ | $0.4 \pm 0.1$ | $5.4 \pm 1.2$ | | $0.74 \pm 0.05$ | 5 |
| | 100 | $0.5 \pm 0.1$ | $0.5 \pm 0.1$ | $7.0 \pm 1.3$ | | $0.38 \pm 0.05$ | 5 |
| | 1000 | $0.06 \pm 0.01$ | $0.9 \pm 0.1$ | $8.8 \pm 1.8$ | $49 \pm 7$ | $0.17 \pm 0.03$ | 5 |

Values are represented as mean ± S.E.M.

### Milliseconds to second open/close kinetics

Upon activation by glycine, heteromeric GlyRs transition from the closed resting state to the open conductive state(s), which in turn transforms to activated but non-conductive desensitized state(s)(23). The transition kinetics between these states directly influence total ion flux in transient physiological processes (such as action potentials) but have yet been systematically characterized. We first determined the kinetics between the closed and open states (Fig. 2). The open/close kinetics of α1β GlyR roughly follows the laws of mass-action but deviates from simple second-order reaction. At lower glycine concentrations, the opening is slower with a larger time constant τ_open_ (∼0.6s at 40 μM glycine), which decreases approximately in proportion to increased glycine concentrations until reaching a minimum of 60 ms likely limited by instrument (Fig. 2A-D). The close rate is fast with τ_close_ ∼ 0.3 s at 40 μM glycine, and showed a non-linear steep slowing down beyond 200 μM glycine to ∼1.1 s. This difference from simple second order reaction is likely related to 2 glycine binding being sufficient and necessary for GlyR activation. Consistent with this, the fraction of open GlyR as calculated from *η* = τ_close_/(τ_close_+τ_open_) very closely follows the dose-response curve obtained from glycine titration (Fig. 2I). These measurements demonstrate that α1β GlyR quickly opens and closes following the binding and dissociation of glycine, which allows fast response to action potentials where glycine concentration is transiently elevated.

**Figure 2.**
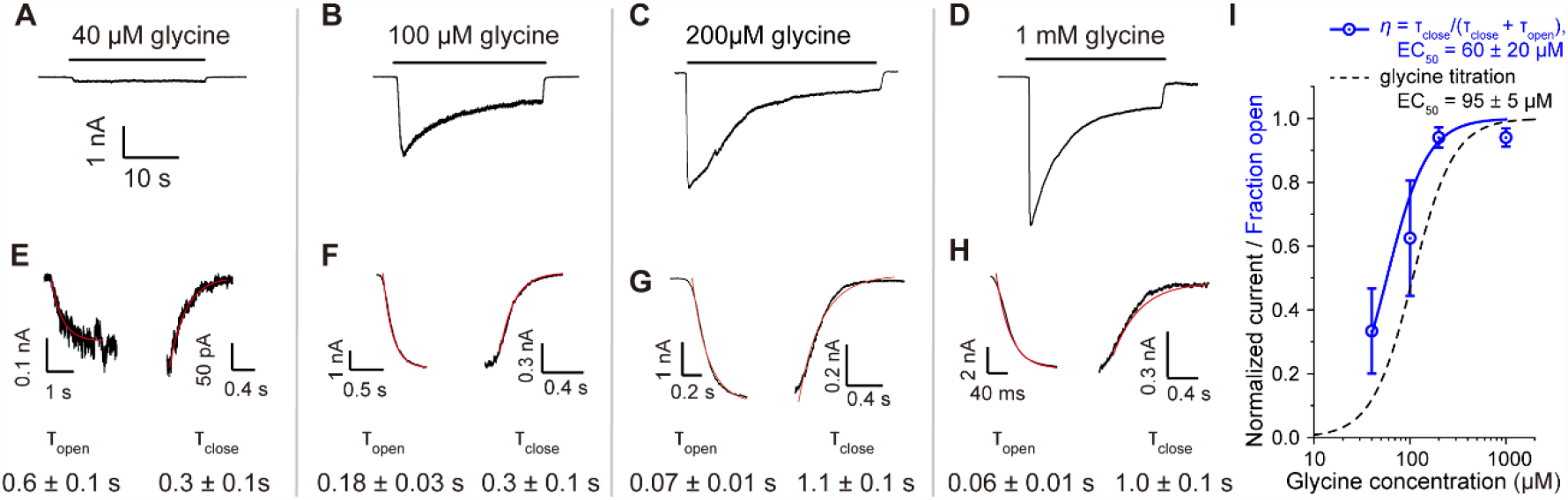
Open and close kinetic measurements of α1β GlyR at varied glycine concentrations. (A-D) Typical process of open, desensitization and close in response to the application and removal of (A) 40 μM, (B) 100 μM, (C) 200 μM and (D) 1000 μM glycine. (E-H) Open and close time courses (n = 5-13 cells) fitted with single-exponential with time constants (mean ± S.E.M.) shown for (E) 40 μM, (F) 100 μM, (G) 200 μM and (H) 1000μM glycine. (I) Fraction GlyR open calculated from open/close rates (blue, with propagated S.E.M.) approximates dose-response measurements (black).

### Desensitization rate is independent of glycine concentration

With sustained application of glycine, α1β GlyR quickly opens, followed by a decrease in conduction owing to transition into non-conductive desensitized state(s). We characterized the transition rate from open to desensitization under varying glycine concentrations (Fig. 3A-D, See Supplementary Fig. 3 for representative complete recordings). Clearly, desensitization followed a single exponential process with comparable time constants (∼ 5-8s) across all tested glycine concentrations spanning 40 μM to 1 mM (∼0.4 – 10x EC_50_). Such invariant kinetics clearly differs from open rate which scales with glycine concentration (Fig. 2), suggesting that open to desensitized state transition is independent of the number bound glycine but an intrinsic property of activated α1β GlyR.

**Figure 3.**
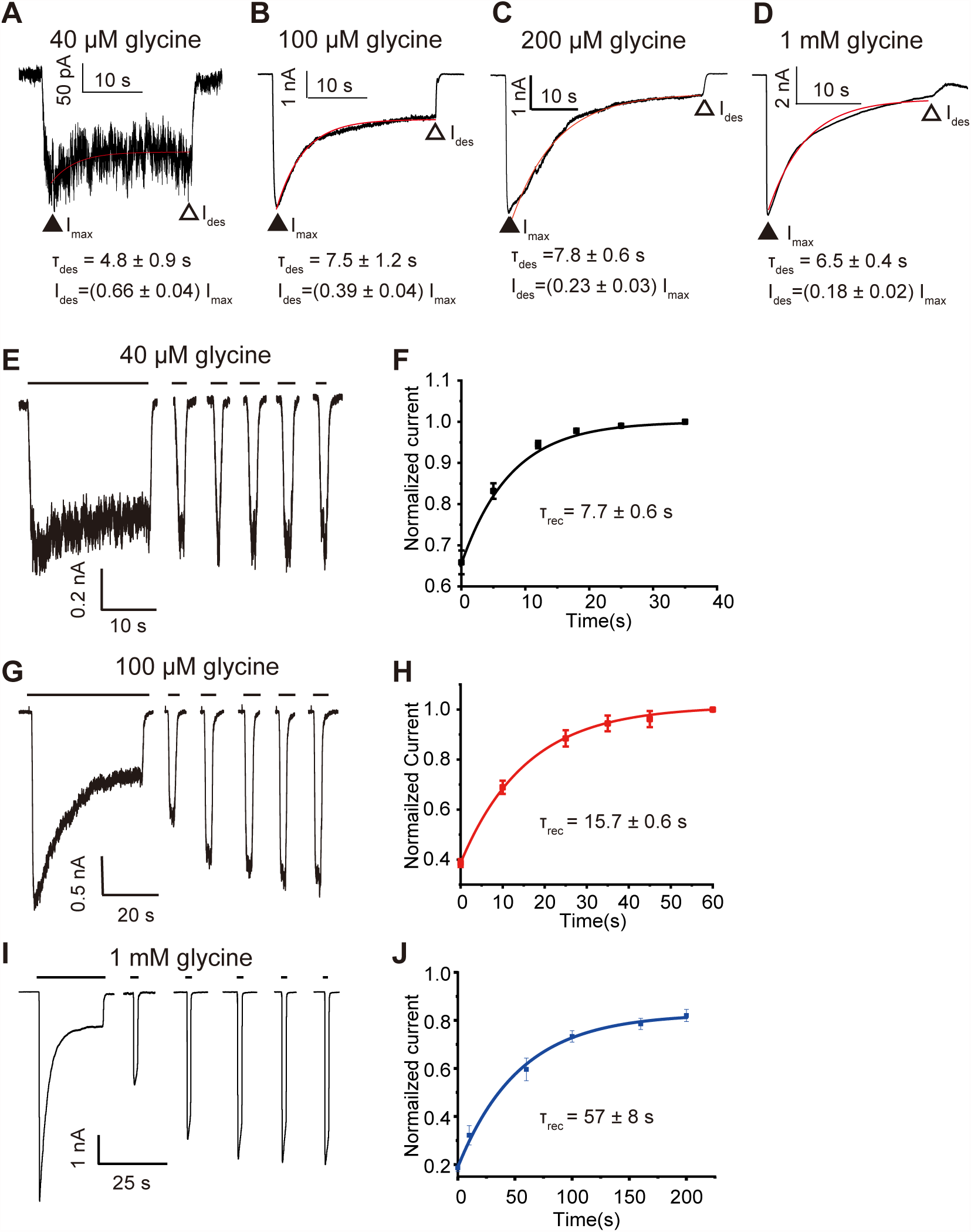
Differential glycine-dependence of desensitization rate, extent, and its recovery. (A-D) GlyR α1β desensitization responses in the presence of glycine (A) 40 μM, (B) 100 μM, (C) 200 μM and (D) 1000 μM are shown. Typical time courses and best exponential fits are shown, with time constants *τ* listed (n = 5 – 13 cells, mean ± S.E.M.). (E, G, I) Representative recordings measuring the recovery after desensitization of GlyR α1β in the presence of (E) 40 μM, (G) 100 μM and (I) 1000 μM glycine. (F, H, J) Averaged desensitization-recovery curves from the sustained application of (F) 40 μM, (H) 100 μM and (J) 1000 μM glycine with exponential fits and time constants shown (n = 6-10 cells, mean ± S.E.M.).

### Higher glycine concentration results in more desensitization and slower recovery

Although the rate of desensitization is unaffected, the extent of desensitization, as well as the recovery rates are directly determined by glycine concentrations. At 40 μM glycine, ∼66% of current remains after desensitization. This number decreases as glycine concentration increases, reaching ∼18% at 1 mM glycine (Fig. 3A-D), a decrease of around 4 folds. Coincident with this, α1β GlyR recovers from desensitization faster when activated by lower glycine concentrations. The recovery time constant τ_rec_ at 40 μM glycine was ∼ 8 s but reached ∼ 57 s at 1 mM glycine (Fig. 3E-J). This ∼ 7 folds increase in desensitization recovery time constant roughly accounts for the ∼ 4 folds increase in the extent of desensitization, as the open to desensitization rate kept constant.

### Intracellular M3-M4 loops do not affect α1β GlyR gating by glycine

The large intracellular M3-M4 loops has been proposed to modulate Cys-loop receptor function in multiple cases, especially in occasions where post-translational modification on these loops clearly affect GlyR function likely under disease situations(5, 8, 24). However, some functional effects are conflicting in specific cases(25-28). We set out to characterize the functional effects of α1 and β M3-M4 loops without any intentional post-translational modifications through truncation (α1_s_) and insertion of irrelevant protein (GFP for β_sg_, Fig. 4A).

**Figure 4.**
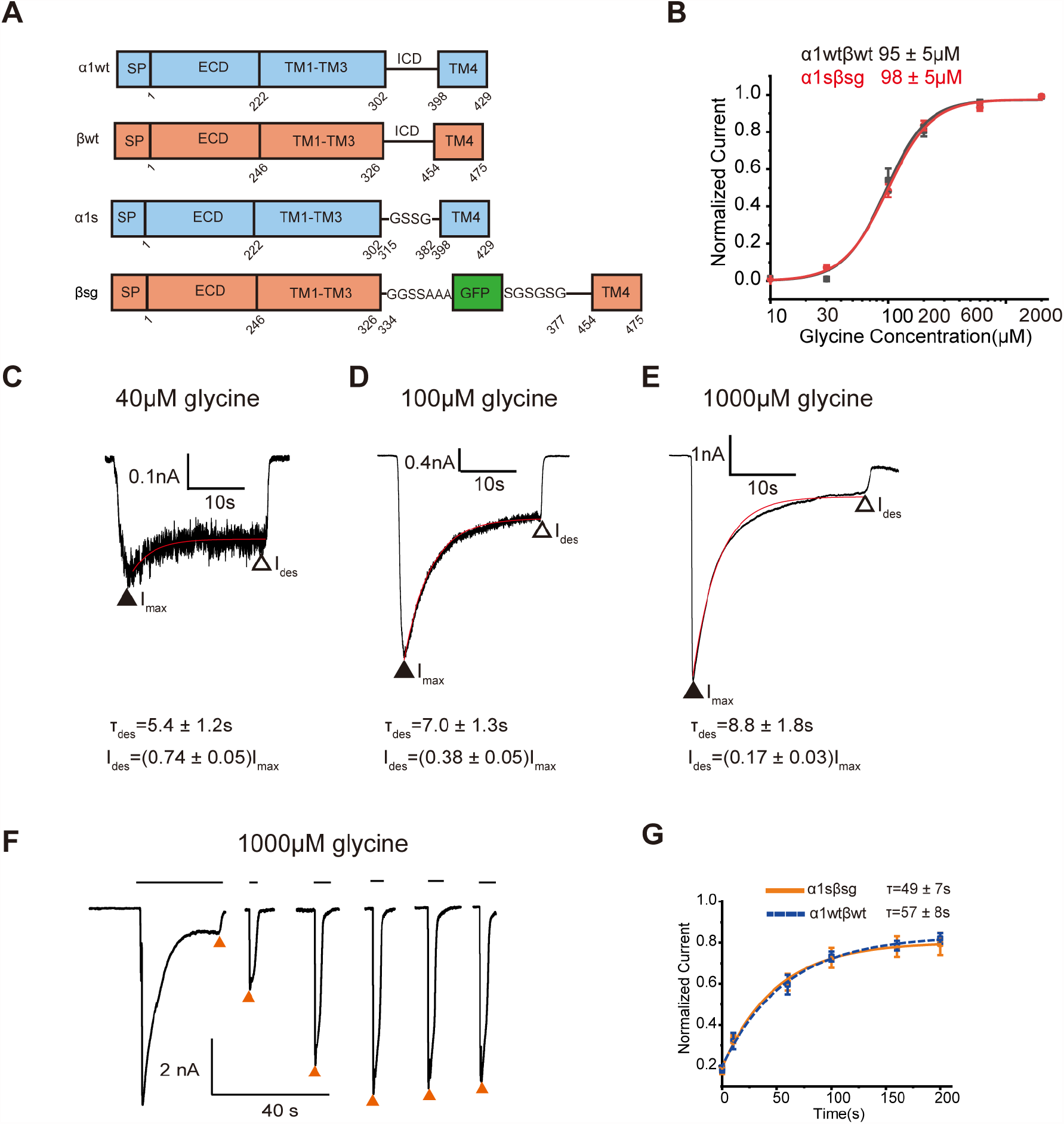
Gating property comparisons between the wild-type and M3-M3 large intracellular domain deleted α1β GlyR. (A) Domain organizations of wildtype (α1wt and βwt) and truncated (α1s and βsg) subunits. SP, signal peptide; ECD, extracellular domain; TM, transmembrane; ICD, intracellular domain; GFP, green fluorescent protein. (B) Glycine response curves of α1wtβwt and α1sβsg (n = 12–17cells, mean ± S.E.M.), normalized to 1 mM glycine in each cell. (C-E) Desensitization rate measurements of truncated α1sβsg GlyR at (C) 40 μM, (D) 100 μM and (E) 1000 μM glycine, with typical exponential fits and time constants shown (n = 6 cells, mean ± S.E.M.). (F) Representative desensitization recovery recording of GlyR α1sβsg from sustained application of 1000 μM glycine. (G) Recovery rate of α1sβsg GlyR fitted to exponential with time constant shown (orange, n = 6-10, mean ± S.E.M.). Wild-type recovery is shown as dashed line for comparison.

Modifications of α1 and β M3-M4 loops did not show appreciable functional effects in the thermodynamics or kinetics of α1β GlyR activation by glycine. The apparent glycine affinities were indistinguishable with EC_50_ ∼ 100 μM (Fig. 4B). In addition, similar invariant rate but glycine concentration-dependent extent during desensitization was observed (Fig. 4C). Both wild-type and mutants had similar slow recovery rate with ∼ 50–60 s time constants when activated with 1 mM glycine (Fig. 4D). Clearly, the modifications we made did not have functional effects on α1β GlyR in glycine activation. These findings suggest that the intracellular loops likely do not intrinsically modulate heteromeric GlyR function. Instead, post-translational modifications on these loops are more relevant.

### A mechanistic model for α1β GlyR gating

The peculiar gating properties of α1β GlyR point to complicated and confounding gating mechanisms. For instance, how does β subunit contribute to gating as its allosteric sites are neither necessary nor essential for activation? How does the same glycine-activated ECD conformation result in distinct open and desensitized states? How does glycine concentration modulate the extent and recovery rate of desensitization without change its forward rate? To address these questions, we propose a gating model that consolidates all the above gating properties and provide a comprehensive description encompassing 6 closed states, 6 open states and 6 desensitized states depending on glycine occupation of the 3 α-α and 2 α-β / β-α sites and kinetics of interchange (Fig. 5A, B).

**Figure 5.**
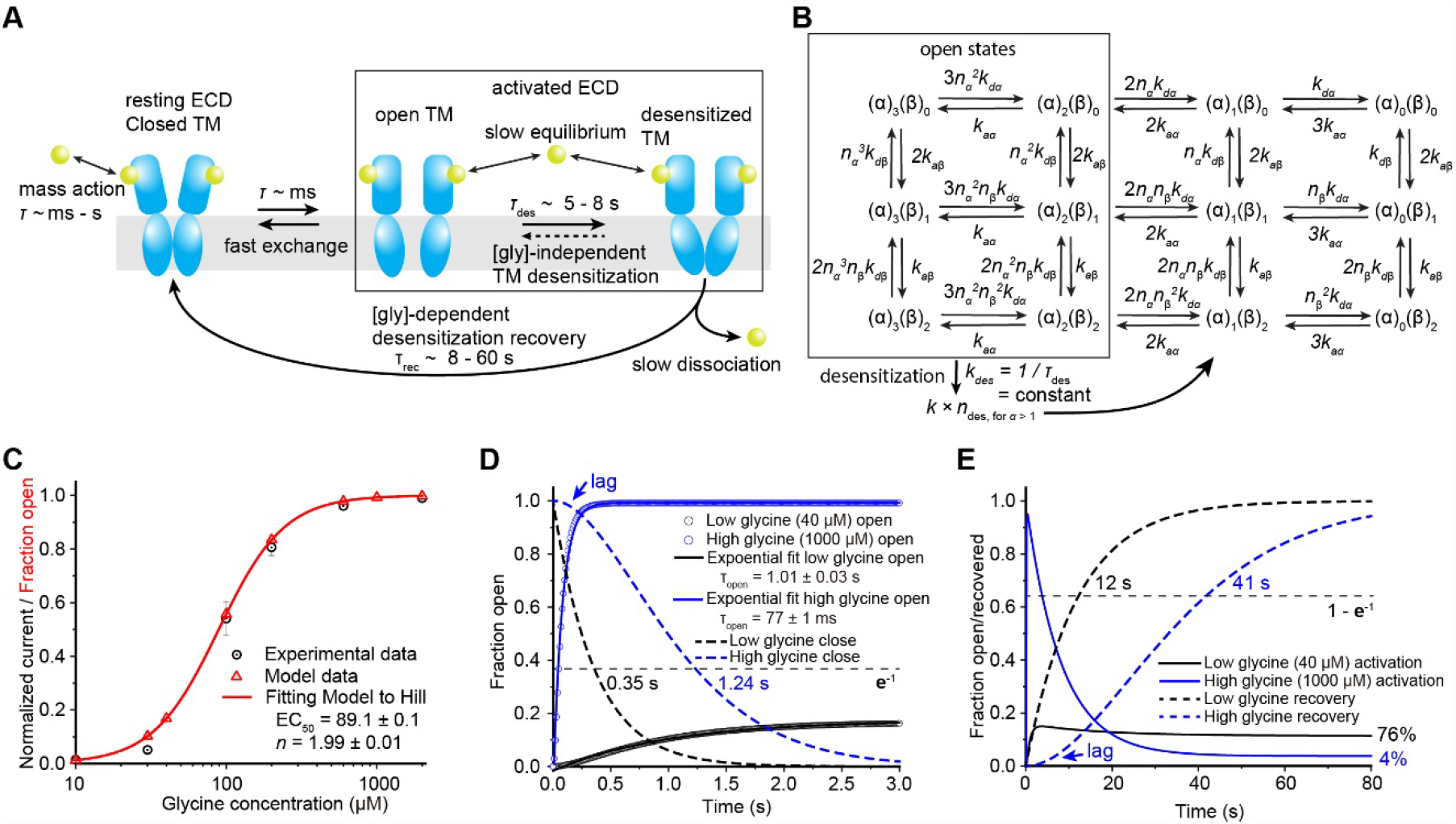
A gating model of the α1β GlyR. (A) Schematic of the gating cycle. (B) Reaction schematic of the gating model, with rate constants shown. Glycine equilibrates quickly at allosteric sites at both α-α (association, dissociation rate constants: *k*_*aα*_, *k*_*dα*_) and α-β (*k*_*aβ*_, *k*_*dβ*_) interfaces with cooperativity index *n*_α_ and *n*_β_ and dominates close-open transitions. ECD activates when 2 or more α-α sites are occupied. Open TM spontaneously relaxes to desensitized TM with an intrinsic constant rate (*k*_des_ = 1 / *τ*_des_), which scales (slows) kinetics by *n*_des_ folds. Please see methods for detailed numbers. (C) 2 or more α-α sites activation closely follows glycine dose-response measurements (reproduced from Fig. 1C). (D) Glycine-dependent open-close kinetics and (E) desensitization rate, extent and recovery rates predicted by the model. Predicted numbers recapitulates experimental ones.

This gating model is based on two general observations currently available of GlyRs (Fig. 5A). First, the ECDs of all 5 subunits are cooperative (thus change simultaneously) and have only two conformations (when gated by glycine), resting and activated, which correspond to the closed and open/desensitized TM conformations, respectively. All available GlyR structures have (pseudo-)5-fold symmetric ECDs that are either in the resting (closed TM) or activated (open/desensitized TM) conformations(14, 15, 29-34). In addition, since glycine is more engulfed in the activated ECD conformation, the binding/dissociation kinetics is likely much slower compared to resting state ECD. Second, the open and closed states are constantly under fast exchange. Single channel recordings show millisecond time scale kinetics between these states(14, 20, 35, 36). This model closely follows α1β GlyR activation properties by glycine. Although glycine binding at 2 α-α sites is necessary and sufficient for activation, abolishing α-β / β-α sites lowered apparent affinity rough by 1.4 folds each (Fig. 1). This is modeled through cooperativity between glycine bindings at the α and β subunits (Fig. 5B, *n*_*a*_ = *n*_*b*_ = 0.7 ≈ 1/1.4). Assuming the same association/dissociation rate constants at α and β subunit interfaces, *k*_aα_ = *k*_aβ_ = 0.011 μM^-1^ and *k*_dα_ = *k*_dβ_ = 2, equilibrium open channel fraction (Fig. 5C red) agrees very well with measurements (Fig. 5C black) on both EC_50_ (89 v.s. 95 ± 5) and apparent Hill slope (2.0 v.s. 2.5 ± 0.4, see Table 1). Glycine association/dissociation with the resting state ECD follow laws of mass action at ∼60 ms to 1 s time scales at physiological glycine concentrations (Fig. 2). Since the open/close states exchange rates are much faster on the millisecond scale, glycine equilibrium rates dominate the close/open kinetics (Fig. 5D) as observed experimentally in Figure 2.

The confounding desensitization properties are also faithfully recapitulated by this gating model. Activation of ECD forces the TM into a meta-stable open state, which then relaxes to a more stable desensitized state (*k*_des_ = 0.12 ≈ 1 / 8 s, see Fig. 3 A-D). Since the open to desensitization transition is a Poisson intrinsic of the TM, the rate is naturally unaffected by glycine concentrations (Fig. 5E, Fig. 3A-D, Fig. 4C-E decay of solid lines). The stable desensitized TM conformation in return locks the ECD in the activated state, from which the dissociation of glycine is much slower (Fig. 5B, rate constants scaled by 1/*n*_des_ = 1/0.03 ≈ 30 folds). This leads to the much longer recovery time from desensitization compared to closing (Fig.5D v.s. E dashed lines, Fig. 2 v.s. Fig. 3E-J and Fig. 4F, G). Higher glycine concentrations occupy more sites, which require longer to dissociate below the activation threshold (2 α-α sites, Fig. 1), leading to a concentration-dependent recovery rate (Fig. 3E-J, Fig. 4F, G and Fig. 5E dashed lines). Slower recovery also results in less available channel for activation, leading to a decreased percentage of residual current after desensitization (Fig. 5E solid lines, Fig. 3A-D, Fig. 4C-E after desensitization).

## Discussion

Through systematic electrophysiology measurements and mathematical modeling, we quantitatively characterized the gating mechanism of α1β GlyR and identified underlying fundamental principles. We show that glycine binds cooperatively to all allosteric sites and activates ECD when 2 or more sites at α-α interfaces are bound. Activated ECD drives the TM into a semi-stable open conformation, which collapses spontaneously into a more stable and kinetically slow desensitized state that is compatible with the same activated ECD conformation. Slow glycine kinetics in the activated ECD conformation leads to the much slower (compared to open-close transition) and glycine concentration-dependent desensitization recovery rate. The contribution of individual allosteric site in GlyR activation has been unclear due to the lack of such information in the 5-fold symmetric homomeric GlyR structures(29-34, 37), and further confounded by the dispute over the α:β stoichiometry(21, 22, 38-42). Recently published heteromeric GlyR structures showed an invariable 4:1 α:β stoichiometry, and a minimal of 2 occupied α-α sites in the open or desensitized states(14-16), consistent with our finding of 2 α-α sites being necessary and sufficient for activation. The necessity of α-α sites is likely related to the asymmetric opening caused by the widening of α-α, but not α-β subunit interfaces(16). In addition, 2 α-α sites out of 5 being sufficient means multiple ligand binding states and routes lead to GlyR activation, consistent with functional observations(35, 37). Coincidentally, structures of other receptors in the Cys-loop family also show a 2-site activation pattern. For instances, in α1β3(19) and α1β3γ2(43) GABA(A) receptors, only 2 GABA are bound at the β-α interfaces, while in nicotinic acetylcholine receptors 2 nicotine binding at the α-β interfaces activates the channel(17). 2-site activation lowers the apparent Hill slope and likely benefits receptor response to lower ligand concentrations.

How ligand binding to the ECD drives GlyRs to both open and desensitized states have been puzzling, given that the ECD conformation in both states are identical. More profound desensitization at higher glycine concentrations seemingly points to a mechanism where partial occupancy leads to open while more causes desensitization. However, this is inconsistent with ECD conformation being the same for both open and desensitized states – how would the TM know which state to be in when the ECD, messenger of ligand binding, is the same? Glycine concentration-independent desensitization rate further suggest that desensitization is a process intrinsic to TM. Activated ECD conformation induces the meta-stable open TM, which spontaneously relaxes to the more stable desensitized state. Desensitized state clearly exchanges much slower with other states as apparent compared to open-close transitions, which detains ECD in the activated conformation until glycine dissociates with much slower kinetics. More glycine binding resulting in slower dissociation, and thus more desensitization and slower recovery.

The quantitative model we proposed here closely follows experimental behavior of α1β GlyR through the full gating cycle. We believe this model reflects underlying basic principles. Of course, due to intrinsic limitations of mutagenesis experiments (for example, mutations usually lower affinities instead of ideally abolish completely), small variations are possible., This model might also reflect generally the gating principles of the Cys-loop receptors, given their similarities both in structures and functional properties, with receptor-specific parameters.

## Materials and Methods

### Glycine receptor constructs

The human glycine receptor α1 (NCBI: NP_001139512.1) and β (NCBI: NP_000815.1) sequence were amplified from cDNA clones (McDermott Center, UT Southwestern Medical center). The human glycine receptor α1 was subcloned in the pEG-Bacmam vector and glycine receptor β was subcloned in the pLVX-IRES-ZsGreen1 vector (Clonetech). We generated human α18Qβ, α18QGGSβ, α18QGGSGGSβ, α18QGGSGGS3Qβ, α118Aaβ, α119Aaβ concatemer construct by connecting human glycine receptor α1 and β (without signal peptide) with QQQQQQQQ, QQQQQQQQGGS,QQQQQQQQGGSGGS, QQQQQQQQGGSGGSQQQ,LHPGSIGGSGGSGRAT and LHPGSIGGSGGSGGSGRAT peptide(figure supplement 1C). The peptide was made by PCR overlap extension. All α1 and β mutants was generated using site-directed mutagenesis. The α1s sequence was derived by substitution of M3/M4 loop (residues R316-P381) by GSSG peptide. For the βsg construct, residues N3344-N377 (M3/M4 loop) was removed from β and GGSSAAA-mEGFP-SGSGSG was inserted. GlyR α1β structure from Liu et al., glycine bound open, PDB ID code 8DN5. The sequence of the reading frame in all constructs was confirmed by Sanger sequencing of the full open frame by Eurofins Genomics.

### HEK293T cell culture and transfection

HEK 293T cells (Purchased from ATCC) were grown at 37 °C, 5% CO_2_ incubator in DMEM (Gibco) containing with 10% (v/v) heat-inactivated fetal bovine serum (Corning), 100 units/ml of penicillin G, 100 μg/ml of streptomycin sulfate (Invitrogen). Cells were passaged after reaching around 90% confluence. The cells were passaged every 2–3 d. For expression, cells were plated on 35mm tissue culture dishes (Scientific Laboratory Supplies) with 0.2M/ml containing 2 ml of DMEM supplemented with 10% (v/v) FBS, and then transfected using Lipofectamine 3000 reagent (Invitrogen) with total 0.8μg plasmids /plate at 1α1:3β ratios coding for the above mentioned GlyRs.

GFP fluorescence from β constructs was used to identify the cells expressing the heteromeric α1β GlyRs. Current recordings were conducted after 17-24h transfected at room temperature.

### Electrophysiology

Glycine-induced currents were recorded in the whole-cell patch clamp configuration. The bath solution contained (in mM): 10 HEPES pH 7.4, 10 KCl, 125 NaCl, 2 MgCl_2_, 1 CaCl_2_ and 10 glucoses. The pipette solution contained (in mM): 10 HEPES pH 7.4, 150 KCl, 5 NaCl, 2 MgCl_2_, 1 CaCl_2_ and 5 EGTA. Experiments with varying external glycine concentrations were conducted. The resistance of borosilicate glass pipettes between 2∼7 MΩ. For current recorded, voltage held at -70 mV and a Digidata 1550B digitizer (Molecular Devices) was connected to an Axopatch 200B amplifier (Molecular Devices). Analog signals were filtered at 1 kHz and subsequently sampled at 20 kHz and stored on a computer running pClamp 10.5 software. For glycine EC_50_ values calculation, Hill1 equation was used to fit the dose-response data and derive the EC_50_ (*k*) and Hill coefficient (*n*). For glycine activation, we used 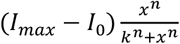, where *I* is current, *I*_*0*_ is the basal current (spontaneous opening current and leak, very close to 0), *I*_*max*_ is the maximum current and *x* is glycine concentration. All start point is fixed at 0 during fit. Measurements were from 7-14 cells, average and S.E.M. values were calculated for each data point. Glycine EC_50_ data analysis was performed by Origin software (Origin Lab). For recovery tau time after desensitization, ExpAssoc1 equation was used to fit the time of GlyR from desensitization state to apo state data and derive the tau time. Measurements were from 5-10 cells, average and S.E.M. values were calculated for each data point. Data analysis of tau time from desensitization to apo state was performed by Origin software (Origin Lab). For tau time calculation of GlyR at open, desensitization and close states, exponential(standard) function was used to fit the trace of open, desensitization and close and calculate tau time. Measurements were from 5-8 cells, average and S.E.M. values were calculated for each data point. Data analysis of tau time was performed by pClamp 10.5 software.

### Gating model and calculations

Differential equation system describing transition between states using rate constants shown in Fig. 5B were resolved numerically with Mathematica (Wolfram Research). Parameters were chosen as follows. Association rate constants for α and β *k*_*aα*_ = *k*_*aβ*_ = 0.011 (μM^-1^); Dissociation rate constants for α and β *k*_*aα*_ = *k*_*aβ*_ = 2; Cooperativity constants for α and β, *n*_α_ = *n*_β_ = 0.7; Desensitization rate constate *k*_des_ = 0.12; Kinetics scaling factor *n*_des_ = 0.03 (meaning a ∼30-fold slow down without shifting equilibrium). Plots were generated using Mathematica and OriginPro (OriginLab).

## Acknowledgments

We thank Robbie Boyed for preparation of tissue culture, and all members of the Wang laboratory for helpful discussions. This work is supported by NIH grant 1R35GM146860 and the McKnight Scholar Award to W.W.

## References

1. A. J. Thompson, H. A. Lester, S. C. Lummis, The structural basis of function in Cys-loop receptors. Q Rev Biophys 43, 449–499 (2010).

2. R. E. Hibbs, E. Gouaux, Principles of activation and permeation in an anion-selective Cys-loop receptor. Nature 474, 54–60 (2011).

3. A. Bode, J. W. Lynch, The impact of human hyperekplexia mutations on glycine receptor structure and function. Molecular brain 7, 2 (2014).

4. W. Xiong et al., Presynaptic glycine receptors as a potential therapeutic target for hyperekplexia disease. Nature neuroscience 17, 232–239 (2014).

5. R. J. Harvey et al., GlyR alpha3: an essential target for spinal PGE2-mediated inflammatory pain sensitization. Science 304, 884–887 (2004).

6. J. W. Lynch, R. J. Callister, Glycine receptors: a new therapeutic target in pain pathways. Current opinion in investigational drugs 7, 48–53 (2006).

7. J. W. Lynch, Y. Zhang, S. Talwar, A. Estrada-Mondragon, Glycine Receptor Drug Discovery. Adv Pharmacol 79, 225–253 (2017).

8. H. U. Zeilhofer, M. A. Acuna, J. Gingras, G. E. Yevenes, Glycine receptors and glycine transporters: targets for novel analgesics? Cell Mol Life Sci 75, 447–465 (2018).

9. H. U. Zeilhofer, K. Werynska, J. Gingras, G. E. Yevenes, Glycine Receptors in Spinal Nociceptive Control-An Update. Biomolecules 11 (2021).

10. V. P. San Martin, A. Sazo, E. Utreras, G. Moraga-Cid, G. E. Yevenes, Glycine Receptor Subtypes and Their Roles in Nociception and Chronic Pain. Frontiers in molecular neuroscience 15, 848642 (2022).

11. T. W. Yu et al., Using whole-exome sequencing to identify inherited causes of autism. Neuron 77, 259–273 (2013).

12. A. Piton et al., Systematic resequencing of X-chromosome synaptic genes in autism spectrum disorder and schizophrenia. Molecular psychiatry 16, 867–880 (2011).

13. J. W. Lynch, Native glycine receptor subtypes and their physiological roles. Neuropharmacology 56, 303–309 (2009).

14. H. Yu, X. C. Bai, W. Wang, Characterization of the subunit composition and structure of adult human glycine receptors. Neuron 109, 2707–2716 e2706 (2021).

15. H. Zhu, E. Gouaux, Architecture and assembly mechanism of native glycine receptors. Nature 10.1038/s41586-021-04022-z (2021).

16. W. W. Xiaofen Liu, Asymmetric gating of a human hetero-pentameric glycine receptor. PREPRINT (Version 1) available at Research Square (2023).

17. A. Gharpure et al., Agonist Selectivity and Ion Permeation in the alpha3beta4 Ganglionic Nicotinic Receptor. Neuron 104, 501–511 e506 (2019).

18. M. M. Rahman et al., Structure of the Native Muscle-type Nicotinic Receptor and Inhibition by Snake Venom Toxins. Neuron 106, 952–962 e955 (2020).

19. V. B. Kasaragod et al., Mechanisms of inhibition and activation of extrasynaptic alphabeta GABA(A) receptors. Nature 602, 529–533 (2022).

20. J. Bormann, N. Rundstrom, H. Betz, D. Langosch, Residues within transmembrane segment M2 determine chloride conductance of glycine receptor homo-and hetero-oligomers. The EMBO journal 12, 3729–3737 (1993).

21. V. Burzomato, P. J. Groot-Kormelink, L. G. Sivilotti, M. Beato, Stoichiometry of recombinant heteromeric glycine receptors revealed by a pore-lining region point mutation. Receptors Channels 9, 353–361 (2003).

22. J. Grudzinska et al., The beta subunit determines the ligand binding properties of synaptic glycine receptors. Neuron 45, 727–739 (2005).

23. J. W. Lynch, Molecular structure and function of the glycine receptor chloride channel. Physiological reviews 84, 1051–1095 (2004).

24. G. Moraga-Cid et al., Modulation of glycine receptor single-channel conductance by intracellular phosphorylation. Scientific reports 10, 4804 (2020).

25. Y. M. Song, L. Y. Huang, Modulation of glycine receptor chloride channels by cAMP-dependent protein kinase in spinal trigeminal neurons. Nature 348, 242–245 (1990).

26. B. Unterer, C. M. Becker, C. Villmann, The importance of TM3-4 loop subdomains for functional reconstitution of glycine receptors by independent domains. The Journal of biological chemistry 287, 39205–39215 (2012).

27. D. Papke, C. Grosman, The role of intracellular linkers in gating and desensitization of human pentameric ligand-gated ion channels. The Journal of neuroscience : the official journal of the Society for Neuroscience 34, 7238–7252 (2014).

28. G. Langlhofer, C. Villmann, The Intracellular Loop of the Glycine Receptor: It’s not all about the Size. Frontiers in molecular neuroscience 9, 41 (2016).

29. J. Du, W. Lu, S. Wu, Y. Cheng, E. Gouaux, Glycine receptor mechanism elucidated by electron cryo-microscopy. Nature 526, 224–229 (2015).

30. X. Huang, H. Chen, K. Michelsen, S. Schneider, P. L. Shaffer, Crystal structure of human glycine receptor-alpha3 bound to antagonist strychnine. Nature 526, 277–280 (2015).

31. G. Moraga-Cid et al., Allosteric and hyperekplexic mutant phenotypes investigated on an alpha1 glycine receptor transmembrane structure. Proceedings of the National Academy of Sciences of the United States of America 112, 2865–2870 (2015).

32. X. Huang, H. Chen, P. L. Shaffer, Crystal Structures of Human GlyRalpha3 Bound to Ivermectin. Structure 25, 945–950 e942 (2017).

33. X. Huang et al., Crystal structures of human glycine receptor alpha3 bound to a novel class of analgesic potentiators. Nature structural & molecular biology 24, 108–113 (2017).

34. A. Kumar et al., Mechanisms of activation and desensitization of full-length glycine receptor in lipid nanodiscs. Nature communications 11, 3752 (2020).

35. R. Lape, A. J. Plested, M. Moroni, D. Colquhoun, L. G. Sivilotti, The alpha1K276E startle disease mutation reveals multiple intermediate states in the gating of glycine receptors. The Journal of neuroscience : the official journal of the Society for Neuroscience 32, 1336–1352 (2012).

36. T. Takahashi, A. Momiyama, K. Hirai, F. Hishinuma, H. Akagi, Functional correlation of fetal and adult forms of glycine receptors with developmental changes in inhibitory synaptic receptor channels. Neuron 9, 1155–1161 (1992).

37. J. Yu et al., Mechanism of gating and partial agonist action in the glycine receptor. Cell 10.1016/j.cell.2021.01.026 (2021).

38. D. Langosch, L. Thomas, H. Betz, Conserved quaternary structure of ligand-gated ion channels: the postsynaptic glycine receptor is a pentamer. Proceedings of the National Academy of Sciences of the United States of America 85, 7394–7398 (1988).

39. T. I. Webb, J. W. Lynch, Molecular pharmacology of the glycine receptor chloride channel. Current pharmaceutical design 13, 2350–2367 (2007).

40. N. Durisic et al., Stoichiometry of the human glycine receptor revealed by direct subunit counting. The Journal of neuroscience : the official journal of the Society for Neuroscience 32, 12915–12920 (2012).

41. Z. Yang, E. Taran, T. I. Webb, J. W. Lynch, Stoichiometry and subunit arrangement of alpha1beta glycine receptors as determined by atomic force microscopy. Biochemistry 51, 5229–5231 (2012).

42. A. Patrizio, M. Renner, R. Pizzarelli, A. Triller, C. G. Specht, Alpha subunit-dependent glycine receptor clustering and regulation of synaptic receptor numbers. Scientific reports 7, 10899 (2017).

43. A. Sente et al., Differential assembly diversifies GABA(A) receptor structures and signalling. Nature 604, 190–194 (2022).

