## Supplemental data for "Gating mechanism of the human α1β GlyR by glycine"

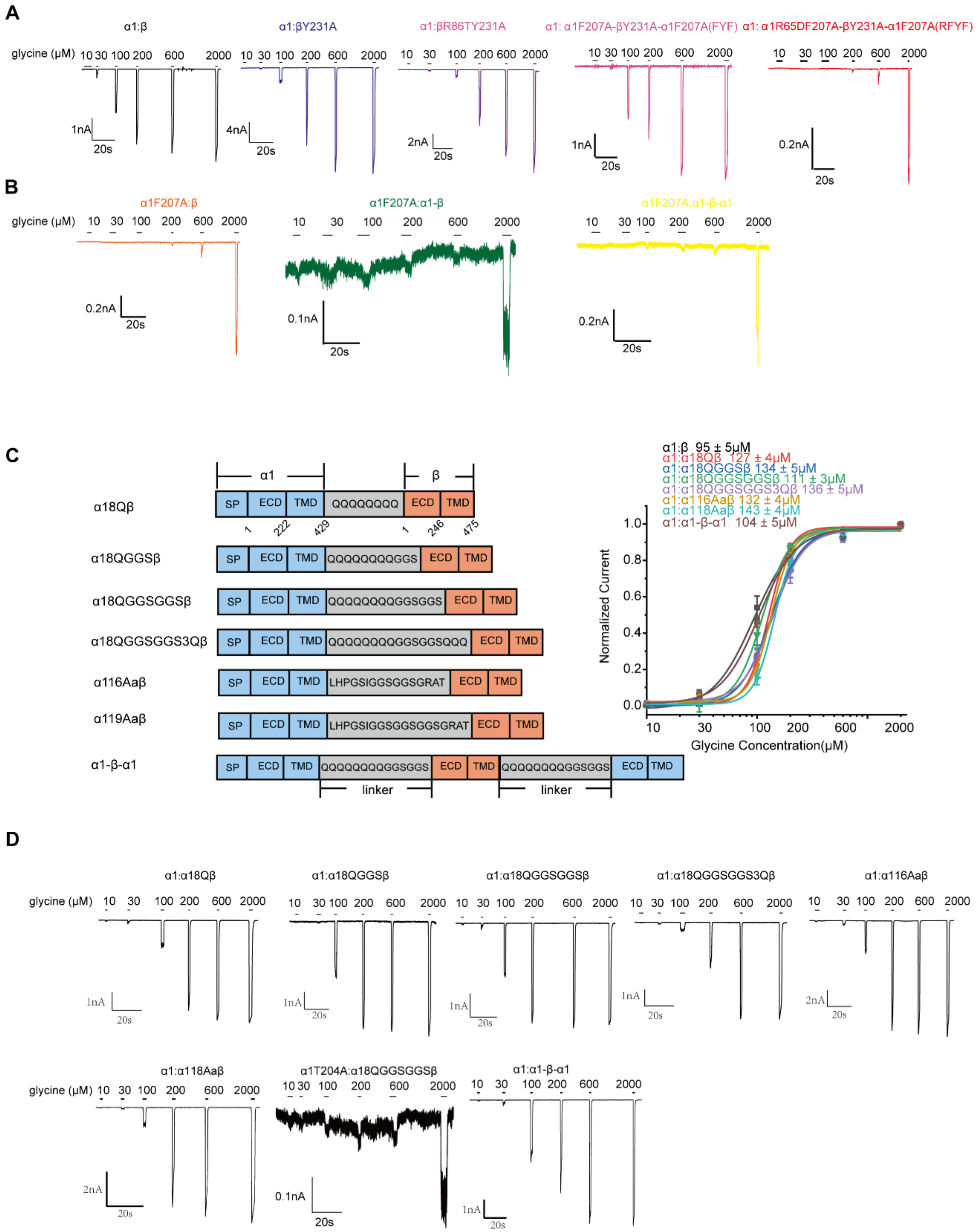


**Supplementary Figure 1 (S1).** Averaged concentration-response curves to glycine of human α1-β and α1-β-α1 concatemers and representative complete whole cell voltage-clamp electrophysiology traces. (A)Typical complete whole cell voltage-clamp electrophysiology traces of human GlyR α1wtβwt and mutants in figure 1C.(B)Typical complete whole cell voltage-clamp electrophysiology traces of human GlyR α1wtβwt and mutants in figure 1E.(C)Left panel, domain organizations of human GlyR α18Qβ, α18QGGSβ, α18QGGSGGSβ, α18QGGSGGS3Qβ, α116Aaβ, α119Aaβ and α1-β-α1 concatemers from up to down. SP, signal peptide; ECD, extracellular domain; TMD, transmembrane domain. Right panel, averaged concentration-response curves to glycine of human GlyR α1wtα18Qβ, α1wtα18QGGSβ, α1wtα18QGGSGGSβ, α1wtα18QGGSGGS3Qβ, α1wtα116Aaβ, α1wtα119Aaβ and α1-β-α1 concatemers. Each curve is constructed from pooling individual concentration-response curves obtained in different cells (n = 8–14, mean ± S.E.M.). Responses are normalized to the response to 1 mM glycine in each cell. (D) Typical complete whole cell voltage-clamp electrophysiology traces of human GlyR concatemers in supplementary figure 1C.

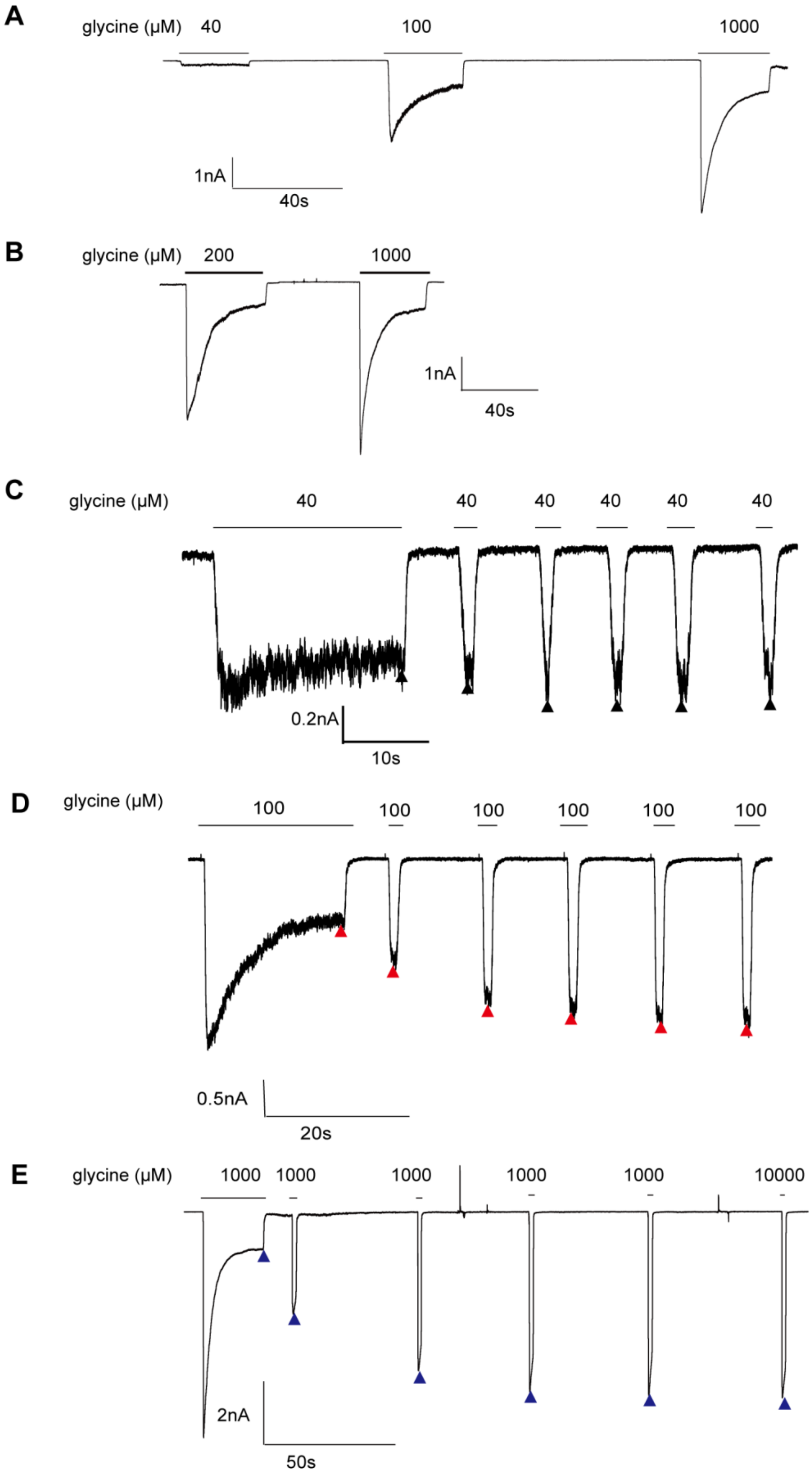
A
glycine (μM) 40
100
1000
1nA
40s
B
glycine (μM) 200
1000
1nA
40s
C
glycine (μM)
40
40
40
40
40
40
0.2nA
10s
D
glycine (μM)
100
100
100
100
100
100
0.5nA
20s
E
glycine (μM)
1000
1000
1000
1000
1000
10000
2nA
50s

**Supplementary Figure 2 (S2).** Typical complete voltage-clamp electrophysiology desensitization and recovery after desensitization traces of human GlyR α1wtβwt under different glycine concentration. (A-B) Typical complete electrophysiology desensitization traces of human GlyR α1wtβwt in presence of 40μM, 100μM, 200μM and 1000μM glycine. (C-E) Typical complete electrophysiology recovery after desensitization of human GlyR α1wtβwt traces in presence of 40μM(C), 100μM (D) and 1000μM(E) glycine.

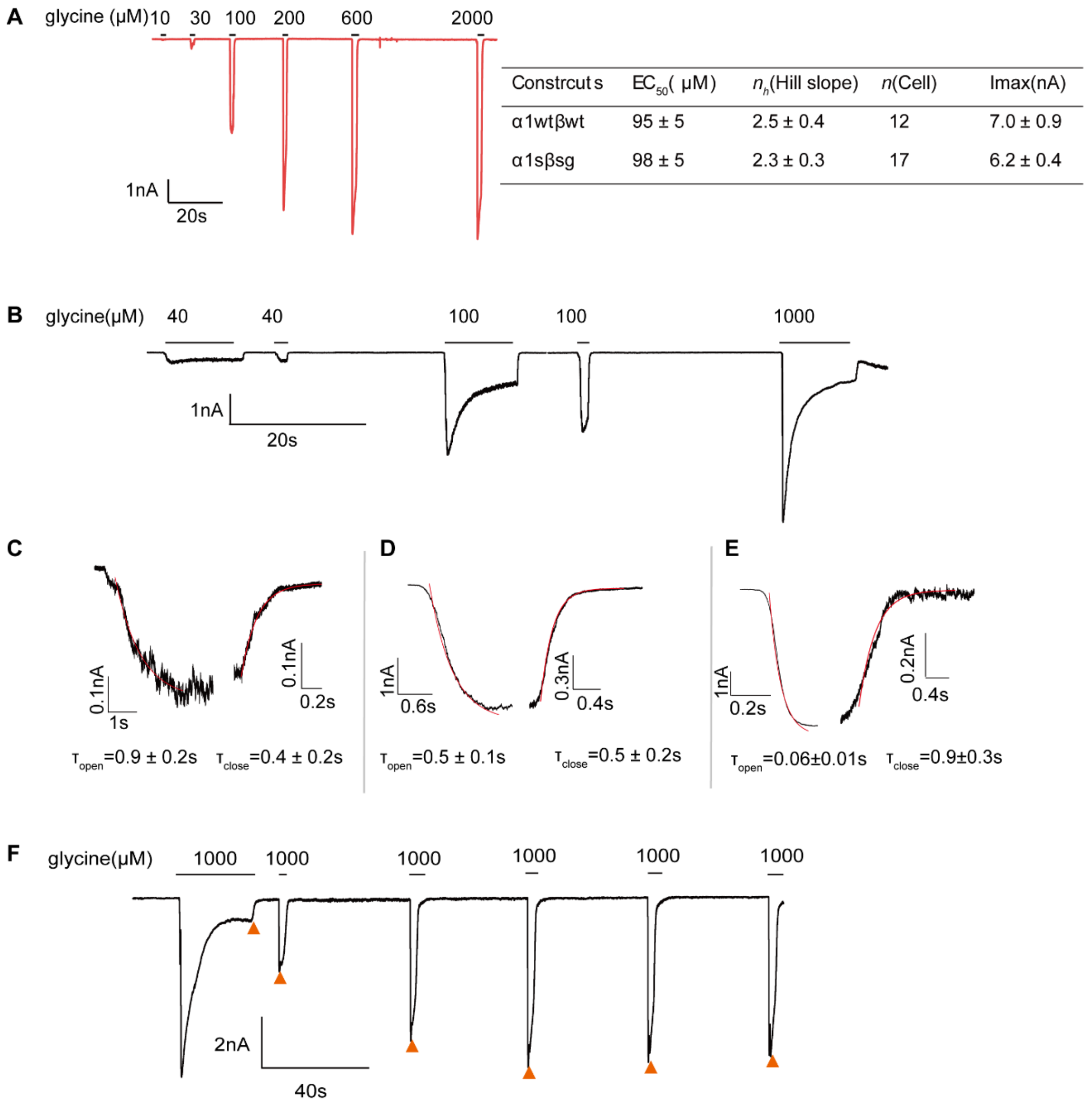


| Constrcuts | $EC_{50}$( μM) | $n_h$(Hill slope) | *n*(Cell) | Imax(nA) |
|---|---|---|---|---|
| α1wtβwt | 95 ± 5 | 2.5 ± 0.4 | 12 | 7.0 ± 0.9 |
| α1sβsg | 98 ± 5 | 2.3 ± 0.3 | 17 | 6.2 ± 0.4 |

**Supplementary Figure 3 (S3).** Typical complete whole cell voltage-clamp electrophysiology traces for human heteromeric GlyR α1sβsg. (A)Left panel, typical complete whole cell voltage-clamp electrophysiology traces of human heteromeric GlyR α1sβsg. $EC_{50}$, $n_h$ (hill slope), n (cell number) and maximum current are listed in right panel. (B)Typical complete electrophysiology desensitization traces of human heteromeric GlyR α1sβsg in presence of 40μM, 100μM and 1000μM glycine. (C-E) GlyR α1sβsg open responses and close responses in the presence of glycine 40μM(C), 100μM (D) and 1000μM(E) are shown. Time constants of best fit are shown in figure E-H. Each tau time is constructed from pooling individual tau time obtained in different cells (n = 6, mean ± S.E.M.). (F) Typical complete electrophysiology recovery after desensitization of human GlyR α1sβsg traces in presence of 1000μM glycine.

**Supplementary Table 1.** Hill fit parameters of glycine dose-response of α1-β and α1-β-α1 concatemers.

| Constrcuts | $EC_{50}$( μM ) | $n_h$(Hill slope) | $n$(Cell) | $I_{max}$(nA) |
|---|---|---|---|---|
| α1:β | 95 ± 5 | 2.5 ± 0.4 | 12 | 7.0 ± 0.9 |
| α1:α18Qβ | 127 ± 4 | 4.4 ± 0.4 | 14 | 6.4 ± 0.7 |
| α1:α18QGGSβ | 134 ± 5 | 3.1 ± 0.3 | 11 | 5.9 ± 0.4 |
| α1:α18QGGSGGSβ | 111 ± 3 | 3.5 ± 0.4 | 12 | 6.2 ± 0.7 |
| α1:α18QGGSGGS3Qβ | 136 ± 5 | 3.0 ± 0.3 | 9 | 3.2 ± 0.6 |
| α1:α116Aaβ | 132 ± 4 | 4.5 ± 0.4 | 9 | 7.0 ± 1.3 |
| α1:α119Aaβ | 143 ± 4 | 4.6 ± 0.4 | 8 | 5.5 ± 0.6 |
| α1:α1-β-α1 | 103 ± 5 | 2.5 ± 0.4 | 10 | 6.2 ± 0.7 |

$EC_{50}$: medium effective concentration of glycine.
$I_{max}$: maximum glycine-induced current amplitude.
